# Sleep Deprivation alters the influence of biological sex on active-phase sleep behavior

**DOI:** 10.1101/2020.02.21.958231

**Authors:** India Nichols, Scott Vincent, September Hesse, J. Christopher Ehlen, Allison Brager, Ketema Paul

## Abstract

Poor sleep is a hazard of daily life that oftentimes cannot be avoided. Gender differences in daily sleep and wake patterns are widely reported; however, it remains unclear how biological sex, which is comprised of genetic and endocrine components, directly influences sleep regulatory processes. In the majority of model systems studied thus far, sex differences in daily sleep amount are predominant during the active (wake) phase of the sleep-wake cycle. The pervasiveness of sex differences in sleep amount throughout phylogeny suggests a strong underlying genetic component. The goal of the current study is to determine if sex differences in active-phase sleep amount are dependent on sex chromosomes in mice.

Sleep was examined in the four-core genotype (FCG) mouse model, whose sex chromosome complement (XY, XX) is independent of sex phenotype (male or female). In this line, sex phenotype is determined by the presence or absence of the *Sry* gene, which is dissociated from the Y chromosome. Polysomnographic sleep recordings were obtained from gonadectomized (GDX) FCG mice to examine spontaneous sleep states and the ability to recover from sleep loss. We report that during the active-phase, the presence of the *Sry* gene accounts for most sex differences during spontaneous sleep; however, during recovery from sleep loss, sex differences in sleep amount are partially driven by sex chromosome complement. These results suggest that genetic factors on the sex chromosomes encode the homeostatic response to sleep loss.

## Introduction

Sleep restriction, a hallmark of life in the modern world, has a substantial negative impact on public health [1]. Compared to men, women report more sleep disturbances such as: fragmented sleep, difficulties falling asleep, restlessness upon awakening, and sleepiness across the daytime [3–6]. Women also experience insomnia more frequently than men, whereas men suffer from sleep apnea more than women [3,4]. Interestingly, in healthy women, quantitative measures of sleep such as polysomnography show that women have better quality sleep than men including more consolidated nighttime sleep, shorter sleep onset latency, and better sleep efficiency [5]. In contrast, men spend more time in lighter stages of sleep at the expense of deeper stages of sleep and have more daytime sleep than women [2]. These studies, among others, provide clear evidence of human gender differences in sleep; furthermore, daytime sleep (napping) in males is observed across the animal kingdom from fruit flies to rodents [7–9]. This conservation of sex differences across species make animal models a useful tool to study these sleep differences.

Sex related peptides and steroids have significant impact on the sleep-wake cycle. In *Drosophila*, transfer of the male sex peptide (SP) during mating induces increased daytime activity [7]. In humans and rodents, estradiol influences sleep by causing more frequent and longer bouts of wakefulness at night and less napping during the day [10,-12]. In rats, estradiol suppresses rapid eye movement (REM) sleep and increases wake during the active-phase [12]. In mice, estradiol reduces non-rapid eye movement (NREM) sleep amount during the active-phase and suppresses the endogenous somnogen prostaglandin D in the ventrolateral pre-optic area (VLPO [13–14]. The role of testosterone in regulating sleep, however, remains elusive. There is lack of a general consensus on the impact of testosterone due to inconsistent reports of its impact on sleep quality [15, 16]. Testosterone increases active-phase NREM sleep in gonadectomized male rodents [13], but other reports indicate that fluctuations in testosterone do not affect sleep-wake architecture [12]. In summary, despite the observed impact of sex hormones and peptides, mouse models indicate that sex-differences in sleep are not exclusively regulated by circulating hormones.

In GDX’d wildtype mice (C57BL/6J) most sex differences are eliminated, but sex differences in the mid-active-phase are still present [17]. This difference is characterized by increased active-phase sleep in males when compared to females. Ehlen et al., 2013 reported that both teste determining factor (*Sry*) and sex chromosome complement contribute to sex differences in sleep during the mid-active phase; this includes spontaneous (during ZT 16-18) recovery sleep (during ZT 18-20) [8]. The effects of Sry and sex chromosome complement on the total 12 hour light phase or 12 hour dark phase, however, was not examined. Therefore, the goal of the present study is to examine the role of sex chromosome complement and testes determining factor, *Sry*, on regulating total active phase sleep.

## Methods

### Animals

The initial breeding pairs of FCG mice on C57BL/6J background were kindly donated by Dr. Arthur Arnold (UCLA, Los Angeles, CA). In this mouse line, the *Sry* gene is expressed on an autosome instead of on the Y chromosome. The resulting phenotypes], include: 1. XY mice with *Sry* and a male urogenital system (XYM); 2. XY mice lacking *Sry* with a female urogenital system (XYF); 3. XX mice with *Sry* and a male urogenital system (XXM); and 4. XX mice lacking *Sry* with a female urogenital system (XXF) [see reference 18].

Mice were maintained at Morehouse School of Medicine under a 12 h-12 h light-dark cycle (LD) in a temperature-controlled vivarium with food and water provided ad libitum. All experiments were performed using the National Institutes of Health Guidelines for the Care and Use of Laboratory Animals and approved by the Morehouse School of Medicine Institutional Animal Care and Use Committee.

### Gonadectomy (GDX)

Mice (8-10 weeks of age) were deeply anesthetized with intraperitoneal injection of ketaset (80 mg/kg) and xylazine (8 mg/kg). In the female mice, an incision was made dorsally through skin and muscle layers slightly above the location of the ovary. The ovary was exposed to the surface and the uterine horn was ligated. Skin and muscle layers were sutured with silk. In male mice, a transverse incision was made at the midline of the scrotum, and fatty and connective tissues were removed to expose the inner sacs that encase the testes. A second incision was made through each casing, and the testes and epididymis were exposed to the surface and ligated. The incisions were sutured with silk.

### Recording Implants

Electroencephalograph (EEG) and electromyograph (EMG) electrodes were implanted in mice immediately after GDX using a prefabricated head mount (Pinnacle Technologies, KS) stabilized by four stainless steel epidural screw electrodes. The first two electrodes (frontal and ground) were located 1.5 mm anterior to bregma and 1.5 mm on either side of the central suture. The second two electrodes (parietal and common reference) were located 2.5 mm posterior to bregma and 1.5 mm on either side of the central suture. Electrical continuity between the screw electrode and head mount was achieved with silver epoxy. EMG activity was monitored using stainless-steel Teflon coated wires inserted bilaterally into the nuchal muscle. The head mount (integrated 2×3 pin grid array) was secured to the skull with dental acrylic. Mice were allowed to recover for at least 14 days before sleep recording.

### EEG/EMG Recordings

One week into surgical recovery, mice were moved to the recording chamber and connected to a lightweight tether attached to a low-resistance commutator mounted over the cage (Pinnacle Technologies, Lawerence, KS). This enabled complete freedom of movement throughout the cage. With the exception of the tether, the home cage environment was maintained. Mice were allowed seven days to acclimate to the tether. Recording of EEG/EMG waveforms began at lights-on on day 8. Data acquisition was performed on a PC running polysomnographic software (Sirenia Acquisition, Pinnacle Technologies, Lawerence, KS). Signals were amplified (10x) and high-pass filtered (0.5 Hz) via a preamplifier. EEG signals were then further amplified, low-pass filtered with a 30 Hz cutoff and collected continuously at a sampling rate of 200 Hz. After collection, EEG/EMG waveforms were classified in 4-sec epochs as: 1) wake (low-voltage, high-frequency EEG; high-amplitude EMG); 2) NREM sleep (high-voltage, mixed-frequency EEG; low-amplitude EMG); or rapid-eye movement (REM) sleep (low-voltage EEG with a predominance of theta activity [6–10Hz]; very low amplitude EMG) by a trained observer. EEG epochs determined to have artifact (interference caused by scratching, movement, eating, or drinking) were excluded from analysis. Artifact comprised less than five percent of all recordings used for analysis.

### Statistics

Graph-Pad Prism 8 was used to create graphs and statistical analysis. Three-way ANOVA was used to determine the effect of phase (light vs dark; within), *Sry* (between), and sex chromosome complement (between) on sleep-wake architecture. Two-way ANOVA was used to determine the effects of *Sry* or sex chromosome complement in dark phase sleep. Tukey Post-Hoc analysis was used to determine differences between each genotype.

## Results

### Differences in spontaneous dark phase sleep amount are dependent on *Sry*

Three-way ANOVA revealed a significant effect of phase on baseline total sleep (F1, 31 = 331.0, P<0.0001), NREM (F1, 31 = 325.5, P<0.0001, and REM (F1, 31 = 246.8, P<0.0001) (figure 1a, 1c, 1e). There was also a significant effect of *Sry* and an interaction between phase and *Sry* in total sleep (F1, 31 = 8.135, P=0.0077; F(1, 31) = 4.939, P=0.0337) and NREM sleep (F1, 31 = 6.843, P=0.0136; F1, 31 = 4.184, P=0.0494). There were no significant differences in the light phase (p>0.05).

**Figure 1.**
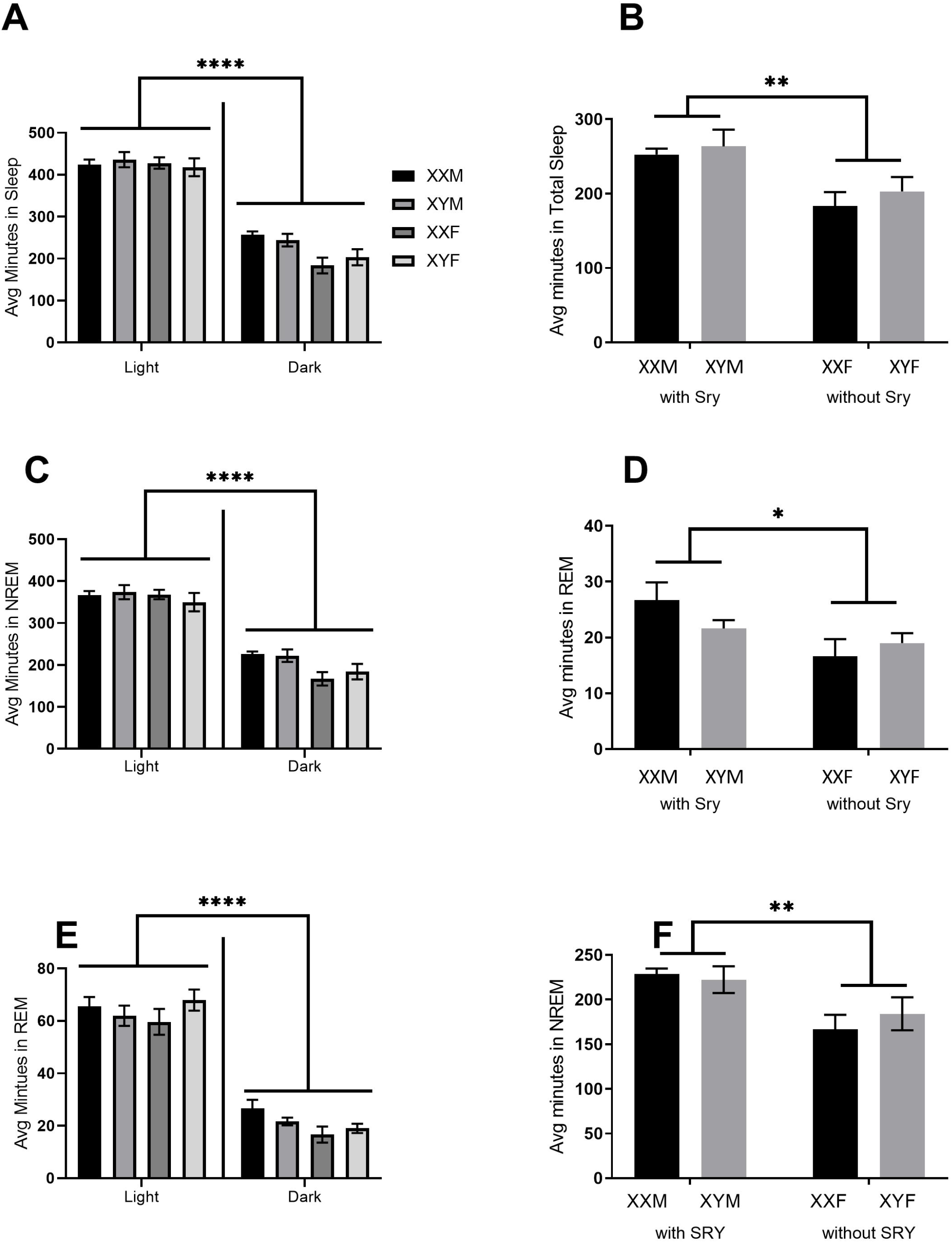
Autosomal transgenic-expression of *Sry* is responsible for sex differences in dark-phase sleep. Repeated measures ANOVA with Tukey post-hoc revealed significant differences in sleep amount during the dark phase. There was a significant effect of phase and all sex differences were observed in the dark phase (A, C, E). In the dark phase, B) mice with *Sry* (XXM, XYM) had more total sleep than mice without *Sry* (XXF, XYF). D) Mice with *Sry* had more NREM and F) REM. Tukey comparison in NREM and REM revealed that XXF mice had less total NREM than XXM and XYM mice, p<0.05 The bars represent the average for 7-10 mice per group and error bars represent standard error. ** denotes significance <0.01 The bars represent the average for 7-10 mice per group and error bars represent standard error. ** denotes significance <0.01

In the dark phase, two-way ANOVA revealed that mice expressing *Sry* (XYM, XXM) had more total sleep (F_1,31_= 14.2; p=.0007), NREM sleep, and REM sleep than mice without *Sry* (XXF, XYF; figure 1b, 1d, 1f). Post hoc analysis revealed that total sleep in XXFs was less than in XYMs (p=0.0147) and XXMs (p= 0.0217; figure 1b). NREM sleep amount was higher in mice with *Sry* (F_1,31_= 11.4, p=.0020; figure 1d) and *Sry* had an effect on REM sleep (F_1,31_ = 5.159; figure 1e), where mice with *Sry* had more REM than mice lacking *Sry*. Fast fourier transformation of EEG waveforms was conducted to determine spectral power of delta (0.5-4.0 Hz) waveforms. There were no significant differences in relative delta between any of the groups (F_3,30_=0.468; p=0.707). There were no effects of sex chromosome complement on any measure of spontaneous sleep amount or NREM delta power.

### *Sry* is responsible for sleep fragmentation in the dark phase

An analysis of sleep fragmentation included number of stage shifts, NREM bouts, brief arousals, and inter-REM interval. Two-way ANOVA showed mice with *Sry* had more bouts of NREM sleep than mice without *Sry* (F_1,25_ =6.591, p=0.0166; figure 2a). It also revealed that mice expressing *Sry* had significantly greater numbers of transitions between sleep and wake (F_1,25_ =9.201, p=0.0056; figure 2b). Two-way ANOVA showed fewer brief arousals (defined as a 4-sec epoch of wakefulness between at least two, 4-sec epochs of sleep) in mice with *Sry* (F_1,25_ = 11.81, 0.0021; Figure 2c).

**Figure 2.**
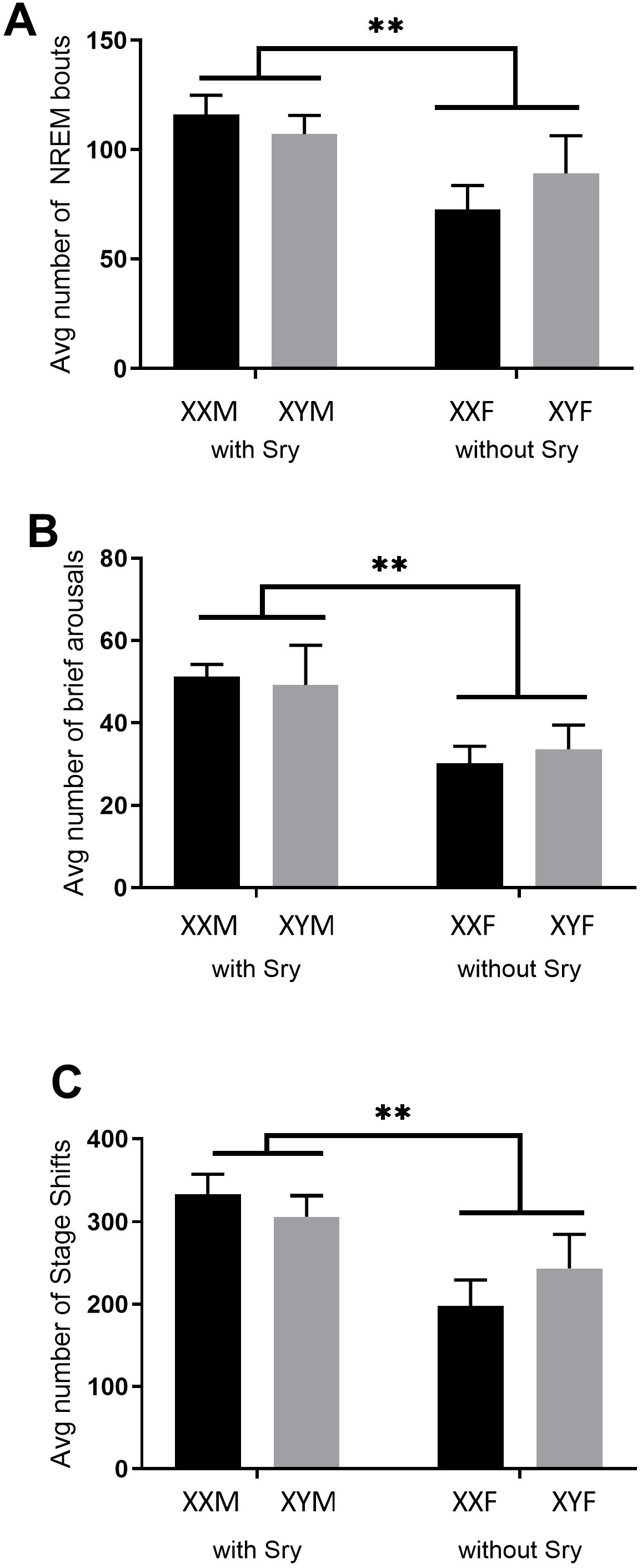
Autosomal transgenic-expression of Sry is responsible for fragmentation in active-phase sleep. Repeated measures ANOVA with Tukey post-hoc comparison revealed significant differences in three measurements of sleep fragmentation. A) Mice with Sry (XXM, XYM) had more NREM bouts than mice without Sry (XXF, XYF). Tukey comparison revealed that XXF mice showed significantly less NREM bouts than XXM and XYM, (p<0.05). B) Mice with Sry (XXM, XYM) had significantly more stage shifts during the active phase than mice without Sry, especially XXF as revealed by Tukey comparison (p<.05). C) Brief arousals were more prevalent in mice with Sry (XXM, XYM) than mice without Sry (XXF, XYF). Tukey comparison revealed significant differences between XXF with XYM and XXM, as well as XYF with XYM, and XXM. The bars represent the average for 7-10 mice per group and error bars represent standard error. ** denotes significance <0.01, * denotes significance <0.05

Sex chromosome complement was not associated with any significant differences in measurements of sleep fragmentation. However, an analysis of inter-REM interval (IRI) showed an effect of sex chromosome complement and *Sry*. IRI is a measurement of the time between two REM episodes, it represents the latency of a full sleep cycle. There was an effect of biological sex (F _3,31_=4.62; p=0.0088) and phase (F _1,31_=50.46; p<0.0001). XXFs have a longer inter-REM interval compared to XYFs (p=0.0142) and mice with *Sry* (XXMs: p<.0001; XYMs: p=0.0171) in the active-phase.

These results suggest that sex *Sry* influences sleep fragmentation in the absence of sex hormones.

### Effects of sex chromosome complement are revealed after sleep loss

In order to determine the role of sex chromosome complement and *Sry* on recovery from sleep loss, FCG mice were subject to 6 hours of sleep deprivation at the beginning of the light phase followed by 18 hours of recovery sleep.

In recovery sleep after 6 hours of forced wakefulness, there was a significant interaction between phase and *Sry* for total sleep amount (F1, 26= 13.48, P=0.0011) and NREM sleep amount (F1, 26 = 14.51, P=0.0008). Phase had an effect on REM sleep amount (F1, 26= 12.83, P=0.0014; figure 3a, 3c, 3e). There was also an interaction between *Sry* and chromosome complement for total sleep (F 1, 26 = 11.76, P=0.0020), NREM (F 1, 26 = 14.51, P=0.0008), and REM (F 1, 26 = 13.08, P=0.0013). There were no differences in the light phase.

**Figure 3.**
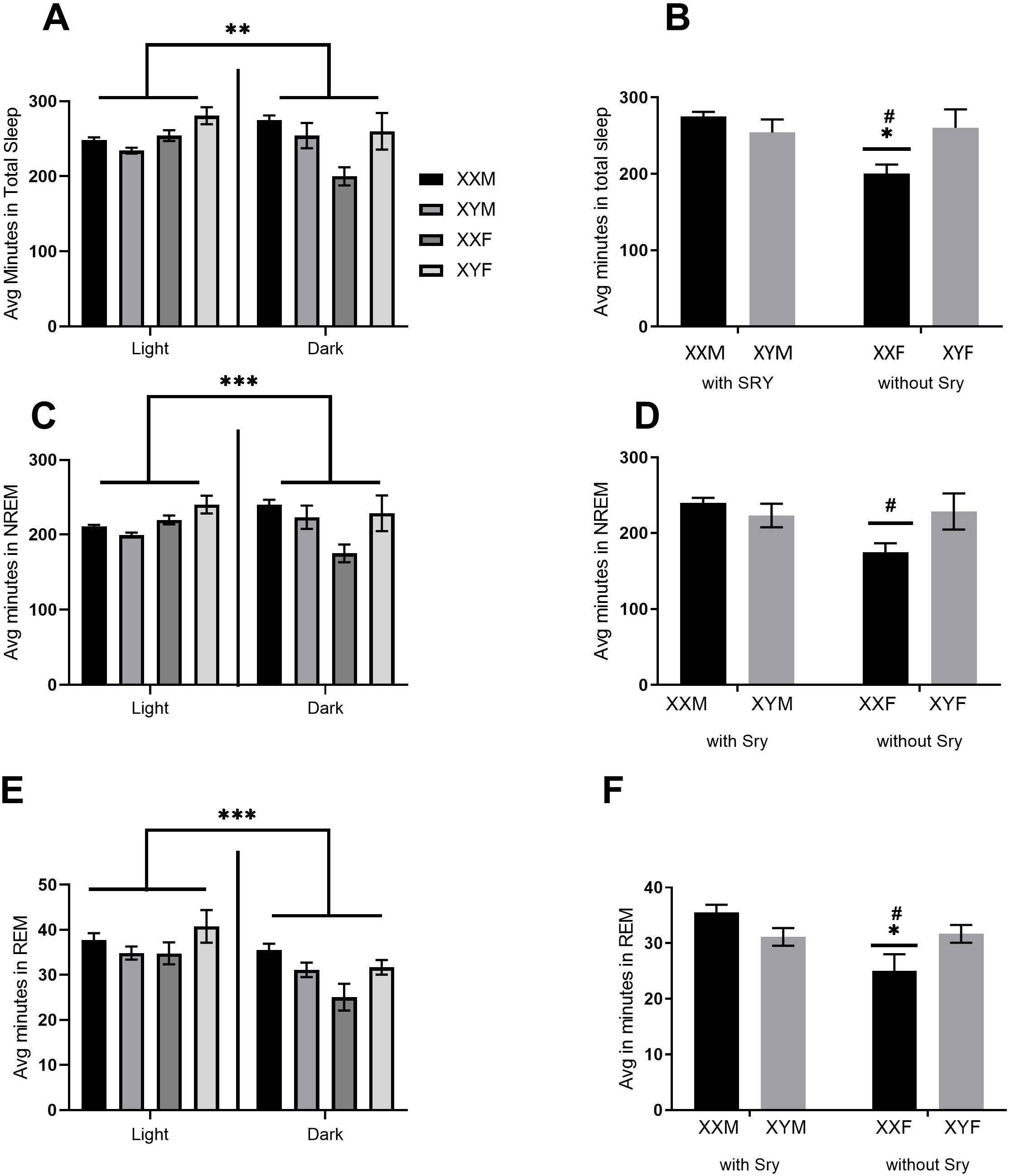
Recovery from sleep loss is dependent on sex chromosome complement. Repeated Measures ANOVA with Tukey Post-Hoc showed significant differences in sleep following 6 hours of sleep loss. There was a significant interaction between phase and *Sry* (A,C,E) where sex differences were observed in the dark phase. B) In total sleep, there was a main effect of *Sry* and an interaction between *Sry* and chromosome complement p<0.05. D) In NREM sleep, there was an interaction between *Sry* and chromosome complement, XXF showed significantly less NREM than mice with *Sry* (XXM, XYM) and Y chromosome (XYF), p<0.05. F) In REM sleep, there was an effect of *Sry* and an interaction between *Sry* and chromosome complement, p<0.05. XXF showed significantly less REM sleep than mice with *Sry* (XXM, XYM) and Y chromosome (XYF), p<0.05. The bars represent the average for 7-10 mice per group and error bars represent standard error. * denotes significance of *Sry* with p <0.05 and # denotes interaction between *Sry* and sex chromosome with p <0.05

Two-way ANOVA showed effect of *Sry* (F_1,26_= 4.905; p=0.0358) and an interaction between *Sry* and sex chromosome complement (F_1,26_ =6.684, p=0.0157 on total sleep (figure 3b. For NREM sleep, there was an interaction between *Sry* and sex chromosome complement (F_1,26_=5.341; p=0.0269) with the effect of *Sry* trending towards significance (F_1,26_ =3.868, p= 0.0600; figure 3d). In REM sleep, there was an effect of *Sry* (F1,26 =5.832, p=0.0231) and an interaction between sex chromosome complement and *Sry* (F1,26=7.221, p=0.0124; figure 3f). FFT of the EEG did not reveal an effect of *Sry* on relative delta power.

Interestingly, sleep deprivation revealed an effect of sex chromosome complement that was not present during undisturbed sleep: XXF mice had less total sleep, NREM, and REM sleep than mice with *Sry* and Y chromosomes (figure 3).

## Discussion

In the present study, we expand upon the findings of Ehlen et al., 2013 to identify the role of sex chromosome complement and *Sry* on sleep during the active-phase sleep, when sex differences are most pronounced [8, 13, 16]. *Sry* effects were apparent under both spontaneous sleep and recovery sleep; however, sex chromosome complement effects were only revealed in sleep recovery. Overall, these findings suggest that the ability to recover from sleep loss is partially regulated by sex chromosome complement.

### Sex differences in spontaneous sleep are regulated by *Sry*

Sex differences during active-phase sleep are well documented. This study reveals that the expression of *Sry* underlies some sex differences in spontaneous sleep. Furthermore, we demonstrate that *Sry* presence results in increased total sleep, REM, and NREM in the active-phase. Both *Sry* genotypes are phenotypic males (XYM & XXM) making these findings consistent with previous reports indicating increased male active-phase sleep before and after GDX [13]. Increased fragmentation in mice with *Sry* present is also consistent with previous findings of less consolidated sleep-wake architecture in males [13, 17]. A significant effect of the Y chromosome on inter-REM interval (decreased in males compared to females) further supports this role in sleep fragmentation [17].

It is important to highlight that a majority of the sex differences we observed were in XX females which have less total sleep, NREM sleep, and REM sleep than all other genotypes; thus, suggesting a role for X chromosome dosage. Therefore, although *Sry* is primarily responsible for sex differences in spontaneous sleep amount, there exists a significant interaction with X chromosome dosage.

### Sex chromosome complement regulates sleep homeostasis

Sleep deprivation altered the influences of sex chromosome complement and *Sry* on sleep-wake architecture during the active-phase. Whereas during undisturbed sleep conditions, *Sry* drove the majority of sex differences, during recovery from sleep loss, sex chromosome complement had a large influence on sleep-wake architecture. These results are intriguing because they may be driven by three separate factors: (a) the presence or absence of *Sry*, (b) the presence or absence of Y chromosome, or (c) X chromosome dosage. Our results suggest all three of these factors interact to drive sex differences in sleep amount.

Future studies in neonatal FCG mice and prepubertal FCG mice may be necessary to reveal potential organizational effects of sex hormones vs sex chromosomes in sleep during development. Since most sex differences are prevalent after puberty, pubertal maturation may permanently alter sleep in females differently than males [19].

## Conclusion

In conclusion, these experiments expand upon previous reports of sex differences in active-phase sleep showing that *Sry* is responsible for sleep amount, sleep fragmentation, and recovery responses and sex chromosome complement is responsible for sleep recovery responses in the absence of hormones. These results show that sex-linked genes play an important role in regulating sleep homeostasis and normal sleep behavior. In a broader context, this data identifies genes and chromosomes that drive sex differences in in sleep.

## Acknowledgements

The authors of this manuscript would like to thank Kyra Clark, Alexandra Hightower, Jeroson Williamson, and Ericka Tummings for their contributions to this research.

## References

1. Armitage R, Hoffmann R, Fitch T, Trivedi M, Rush AJ. Temporal characteristics of delta activity during NREM sleep in depressed outpatients and healthy adults: group and sex effects. Sleep. 2000;23(5):607–617.

2. Kobayashi, R., Kohsaka, M., Fukuda, N., Honma, H., Sakakibara, S., & Koyama, T. Gender differences in the sleep of middle-aged individuals. Psychiatry And Clinical Neurosciences. 1998; 52(2), 186–187. doi: 10.1111/j.1440-1819.1998.tb01021.x

3. Ye L, Pien GW, Weaver TE. Gender differences in the clinical manifestation of obstructive sleep apnea. Sleep Med. 2009;10(10):1075–1084.

4. Suh, S., Cho, N., & Zhang, J. Sex Differences in Insomnia: from Epidemiology and Etiology to Intervention. Current Psychiatry Reports. 2018; 20(9). doi: 10.1007/s11920-018-0940-9

5. Choi-Kwon, S., & Jeon, B. (2017). Are there any differences in subjective and objective sleep measurement?. Sleep Medicine. 2017; 40, e65

6. Suarez EC. Self-reported symptoms of sleep disturbance and inflammation, coagulation, insulin resistance and psychosocial distress: evidence for gender disparity. Brain Behav Immun. 2008;22(6):960–968.

7. Ferguson, C., O’Neill, T., Audsley, N., & Isaac, R. The sexually dimorphic behavior of adult Drosophila suzukii: elevated female locomotor activity and loss of siesta is a post-mating response. Journal Of Experimental Biology. 2015; 218(23), 3855–3861.

8. Khericha, M., Kolenchery, J., & Tauber, E. Neural and non-neural contributions to sexual dimorphism of mid-day sleep in Drosophila melanogaster: a pilot study. Physiological Entomology. 2016; 41(4), 327–334. doi: 10.1111/phen.12134

9. Ehlen JC, Hesse S, Pinckney L, Paul KN. Sex chromosomes regulate nighttime sleep propensity during recovery from sleep loss in mice. PLoS One. 2013; 8(5):e62205.

10. Manber R, Bootzin RR. Sleep and the menstrual cycle. Health Psychol. 1997;16(3):209–214.

11. Moline ML, Broch L, Zak R, Gross V. Sleep in women across the life cycle from adulthood through menopause. Sleep Med Rev. 2003;7(2):155–177.

12. Cusmano DM, Hadjimarkou MM, Mong JA. Gonadal steroid modulation of sleep and wakefulness in male and female rats is sexually differentiated and neonatally organized by steroid exposure. Endocrinology. 2014;155(1):204–214.

13. Paul KN, Laposky AD, Turek FW. Reproductive hormone replacement alters sleep in mice. Neurosci Lett. 2009;463(3):239–243.

14. Mong JA, Devidze N, Goodwillie A, Pfaff, DW. Reduction of lipocalin-type prostaglandin D synthase in the preoptic area of female mice mimics estradiol effects on arousal and sex behavior. PNAS. 2003; 100(25), 15206–15211.

15. Barrett-Connor, E., Dam, T., Stone, K., Harrison, S., Redline, S., & Orwoll, E. The Association of Testosterone Levels with Overall Sleep Quality, Sleep Architecture, and Sleep-Disordered Breathing. The Journal Of Clinical Endocrinology & Metabolism. 2008; 93(7), 2602–2609.

16. Liu, P., Yee, B., Wishart, S., Jimenez, M., Jung, D., Grunstein, R., & Handelsman, D. The Short-Term Effects of High-Dose Testosterone on Sleep, Breathing, and Function in Older Men. The Journal Of Clinical Endocrinology & Metabolism. 2003; 88(8), 3605–3613. doi: 10.1210/jc.2003-030236

17. Paul KN, Dugovic C, Turek FW, Laposky AD. Diurnal sex differences in the sleep-wake cycle of mice are dependent on gonadal function. Sleep. 2006; 29(9):1211–1223.

18. Arnold AP, Chen X. What does the “four core genotypes” mouse model tell us about sex differences in the brain and other tissues? Front Neuroendocrinol. 2009; 30(1):1–9.

19. Zhang J, Chan NY, Lam SP, Li SX, Liu Y, Chan JW, Kong AP, Ma RC, Chan KC, Li AM, Wing YK. Emergence of Sex Differences in Insomnia Symptoms in Adolescents: A Large-Scale School-Based Study. Sleep. 2016; 39(8):1563–1570

